# The human-specific gene *FRMPD2B* couples postnatal experience to the expansion of cortical connectivity

**DOI:** 10.64898/2026.10.30.763086

**Authors:** Eirini Christopoulou, Joseph Zamith, Pauline Bonadonna, Mariana Rivero Romano, Marie Vaudin, Wendy Nguyen, Clarisse Brunet, Corinne Blugeon, Sophie Lemoine, François Giudicelli, Alexandre Baffet, Cécile Charrier

**Author notes:** **Correspondence and requests for materials** should be addressed to Cécile Charrier. These authors contributed equally (Eirini Christopoulou, Joseph Zamith).

## Abstract

The human neocortex is characterized by its uniquely prolonged postnatal developmental period during which experience sculpts neuronal circuits, a process essential to human cognitive abilities. Yet whether human-specific genetic innovations have shaped the activity-dependent mechanisms regulating circuit assembly remains unknown. Here, we identify the human-specific gene *FRMPD2B* as an activity- and experience-regulated modifier of cortical connectivity. *FRMPD2B* is primarily expressed in cortico-cortical projection neurons postnatally and throughout childhood. It promotes dendritic, axonal and synaptic development, and is required for proper morphogenesis in xenotransplanted human cortical neurons. *FRMPD2B* expression is bidirectionally regulated by neuronal activity and experience. This activity-responsiveness was gained after *FRMPD2B* arose through partial gene duplication, when a retrotransposon insertion remodeled its promoter. Our findings highlight how a series of genetic innovations have led to the emergence of a human-specific activity-regulated developmental program through which postnatal experience drives cortical connectivity expansion, linking genome evolution to human brain plasticity.

## INTRODUCTION

The human neocortex is characterized by distinct cellular, molecular and developmental features that emerged during evolution and underlie our capacity for language, social cognition, abstract reasoning, and cumulative culture^1,2^. In particular, human cortical pyramidal neurons (CPNs) develop and mature over a uniquely prolonged postnatal period extending until early adulthood^3– 8^. The first years of life correspond to a phase of intense brain growth and cortical surface expansion, during which human CPNs elaborate complex dendrites and establish the vast majority of their synaptic connections^4,5,9^. By comparison, cortical synaptogenesis peaks during a 4-month period centered around birth in macaques^9^ and most synapses form within three weeks after birth in mice. Ultimately, human CPNs develop longer, more elaborate dendrites and a higher density of dendritic spines than neurons in other great apes or rodents^10–14^. This greater morphological complexity is associated with increased synaptic connectivity, and together with specialized functional properties, enhanced computational power^3,10,15,16^. Combined with the higher proportion of upper layer CPNs in humans, it also supports the expansion of human cortico-cortical connectivity^8^, which provides the physical substrate for the associative processing underlying higher cognition.

The slow, protracted, and mainly postnatal nature of synaptic development in the human neocortex underlies synaptic neoteny and extends the period during which experiences sculpt cortical circuits and neuronal functional properties^3,17–19^. This window of maturation and high plasticity is thought to be critical for human brain development and its disruption is a hallmark of neurodevelopmental disorders such as autism spectrum disorders (ASD) and intellectual disabilities (ID), which frequently combine altered circuit connectivity with impaired processing of cognitive or social experiences^3,20–23^. Yet whether and how human-specific genetic innovations have introduced new activity-dependent mechanisms governing the assembly of cortical circuits remains unknown. Experience-induced changes in neuronal activity induce the expression of activity-regulated genes, which regulate neuronal maturation, wiring and plasticity^19,24–27^. In vitro assays have identified several genes whose activity-dependent regulation in neurons differs between human and other species, reflecting divergence in gene regulatory sequences^28–31^. However, whether these differences contribute to distinct morpho-physiological features of human CPNs is still elusive. Meanwhile, large segmental duplications, one of the main drivers of human genome evolution, have generated human-specific genes regulating the pool of cortical progenitors (ARHGAP11B, NOTCH2NL) or controlling the timing of synaptic maturation (SRGAP2B/C)^8,32,33^, but these genes have so far only been implicated in the regulation of intrinsic, activity-independent mechanisms.

Here, we characterize the human-specific gene *FRMPD2B* (FERM and PDZ Domain Containing 2B) as an activity and experience-regulated gene. *FRMPD2B* arose in the human genome through the partial duplication of the long ancestral gene *FRMPD2A* approximately 2.3 million years ago, at a time coinciding with the emergence of the genus *Homo. FRMPD2B* is fixed in the human genome in the 10q11.22 chromosomal region, whose deletions and duplications lead to a syndrome associated with ID, ASD, absent speech and epilepsy^34^. It presents a unique expression profile among human-specific duplicated genes with primary expression in the human neocortex^35^. Using a combination of in vivo experiments in mouse and xenotransplantated human CPNs, together with functional genomic analyses, we demonstrate that the human-specific gene *FRMPD2B* regulates dendritic, axonal and synaptic development, and is ultimately required for human upper layer CPNs to acquire complex dendritic morphology. We show that *FRMPD2B* expression is bidirectionally regulated by neuronal activity and experience. Its activity-dependent regulation arose after the duplication, when a retrotransposon remodeled its promoter architecture and introduced binding sites for MEF2 transcription factors. These results reveal how a succession of human-specific genetic innovations, a partial gene duplication followed by retrotransposon-mediated promoter remodeling, have led to the emergence of a novel activity-regulated developmental program that contributes to the expansion of cortical connectivity.

## RESULTS

### FRMPD2B is expressed during human cortical neuron morphogenesis

*FRMPD2B* is the product of the partial duplication of the ancestral gene *FRMPD2A*. The 5’ breakpoint of the duplication is located within intron 20 of *FRMPD2A*, and the human-specific copy *FRMPD2B* encompasses the last nine exons of *the ancestral gene* (Fig. 1a)^35^. *FRMPD2A* encodes a 1,309-amino-acid protein containing an N-terminal KIND domain followed by a FERM domain, three PDZ domains and a disordered C-terminal sequence (Fig. 1a). *FRMPD2B* is currently misannotated as a pseudogene in the human reference genome (Refseq: NR_033172.1). Using long-read nanopore sequencing, we found that it encodes a 320-amino-acid protein corresponding to the C-terminal part of FRMPD2A starting from the middle of the second PDZ domain (Fig. 1b). Remarkably, long *FRMPD2A* transcripts were not detected in the human cortex (Fig. 1b). Exon expression analysis of the *FRMPD2* gene family in the human frontal cortex using the GTEx dataset^36^ yielded similar results (Fig. 1c). Therefore, FRMPD2B is not co-expressed with its long parental copy in the cortex and operates independently. Further investigation by leveraging publicly available datasets^37,38^ revealed that *FRMPD2* is selectively expressed in cortico-cortical projecting excitatory neurons (Fig. 1d). Its expression begins in the last weeks of gestation, peaks between 1 and 2 years after birth, remains sustained throughout childhood and then gradually declines in adults (Fig. 1e). In contrast, *FRMPD2* transcripts were barely detected in macaques across all developmental stages, suggesting that *FRMPD2* neocortical expression is a human-specific feature primarily linked to *FRMPD2B*. Together, these results indicate that *FRMPD2B* is predominantly expressed during the period in which human CPNs develop their dendrites and axon and form the majority of their synaptic connections.

**Fig. 1.**
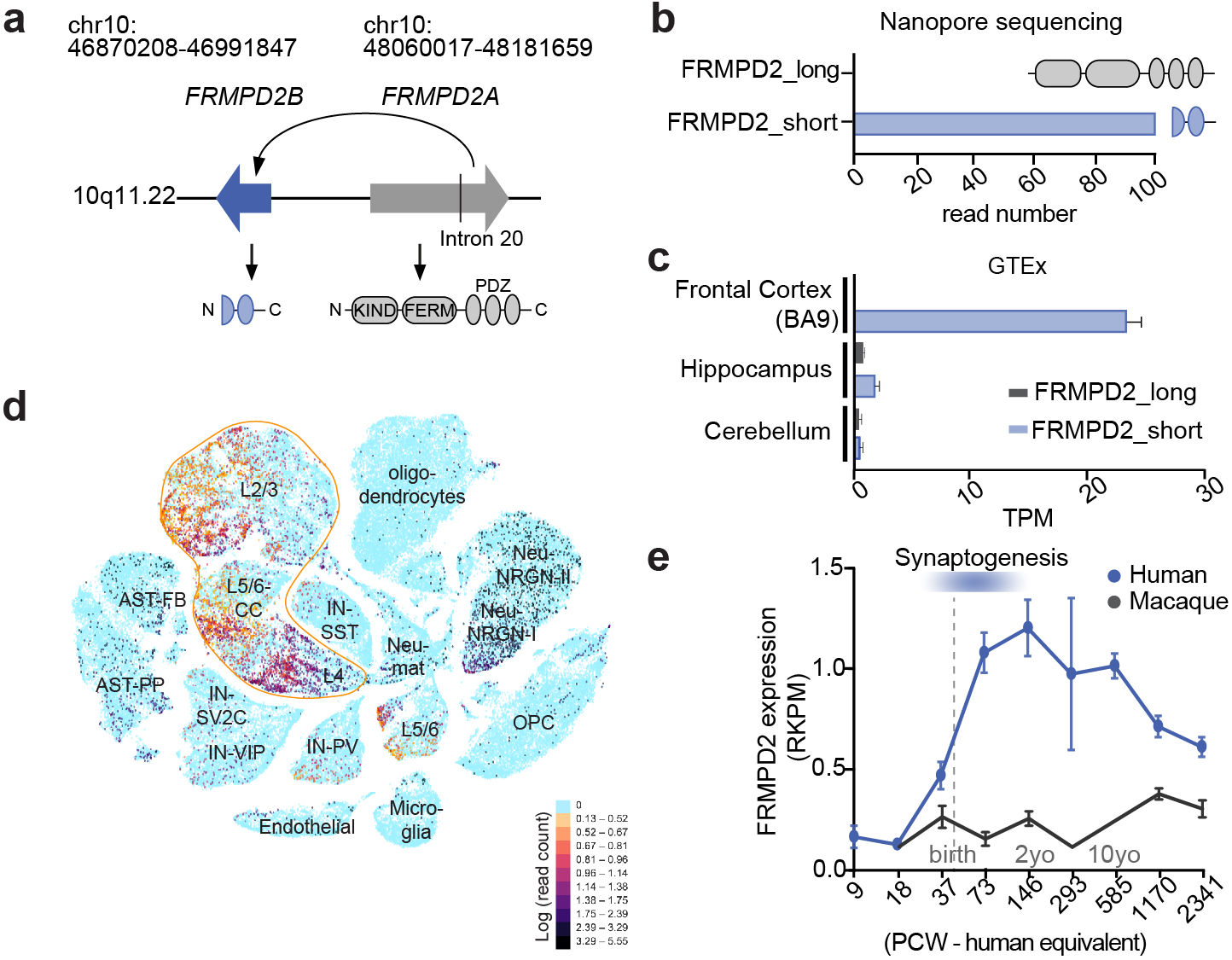
*FRMPD2* human-specific duplication and expression profile in the human neocortex. **a**, Schematic representation of *FRMPD2* human-specific duplication. Top: The duplication that generated *FRMPD2B* occurred within chromosome 10q11.22 and encompasses the last nine exons of *FRMPD2A*. Bottom: Functional domain architecture of the long ancestral protein FRMPD2A and short duplicated protein FRMPD2B. **b**, Quantification of the amount of short and long *FRMPD2* transcripts in the human cortex using nanopore sequencing. **c**, Comparative expression of short and long *FRMPD2* transcripts in the frontal cortex (BA9), hippocampus and cerebellum in the GTEx dataset. TPM: transcripts per million. **d**, t-SNE plot showing *FRMPD2* expression profile across cell types in the human prefrontal and anterior cingulate cortex using single-nuclei RNA-seq data were from^38^ (https://autism.cells.ucsc.edu). The main cell populations expressing *FRMPD2* transcripts (L2-3, L4 and cortico-cortical projecting (CC) L5/6 excitatory neurons) are outlined. **e**, Temporal expression profile of *FRMPD2* from gestational week (GW) 9 to adulthood in humans (blue) and macaques (black) in RNA-seq data from^37^ (http://evolution.psychencode.org). Time is represented in human age equivalent (Post Conception Weeks (PCW)). RPKM: Reads per kilo base per million mapped reads.

### FRMPD2B promotes dendritic, axonal and synaptic development

To determine the role of FRMPD2B in the development of dendrites and synapses, we introduced it along with a cytosolic fluorescent protein (mVenus) in mouse L2/3 CPNs of the somatosensory cortex using targeted in utero electroporation (IUE) (Fig. 2a). To achieve single cell resolution for dendrite reconstruction, we performed ultra-sparse neuronal labelling by co-electroporating a Cre-dependent FLEX-mVenus with low amounts of Cre recombinase (Fig. 2b). In juvenile mice (postnatal day (P)21), FRMPD2B increased the branching and length of basal dendrites without modifying apical dendrites (Fig. 2c, d). FRMPD2B also increased spine density across the dendritic tree (Fig. 2e,f), and dendritic spines in oblique apical dendrites had smaller necks than in control neurons (Fig. S1), suggesting better electrical and biochemical coupling with the dendrite^39–41^. Ultimately, FRMPD2B expression led to a strong increase in mEPSC frequency compared to non-electroporated neurons without modifying mEPSC amplitude or intrinsic membrane properties (Fig. 2i, j and Fig. S2, 3). We then assessed the role of FRMPD2B in axonal development. In vivo, the axons of L2/3 CPNs branch in ipsilateral L2/3 and L5, before crossing the midline through the corpus callosum and branching again in the contralateral side^42^. In ipsilateral L5, FRMPD2B increased axonal branching at both P6 and P21 (Fig. 2k, l). At P21, FRMPD2B also increased the density of presynaptic boutons visualized using GFP-tagged synaptophysin in both L2/3 and L5 (Fig. 2k, l). Consistent results were obtained in cultured cortical neurons. After 5 days in vitro, layer 2/3 CPNs expressing FRMPD2B had a longer and more branched axon than control neurons (Fig. S4). Therefore, FRMPD2B-expressing neurons receive more inputs and establish a greater number of synapses with their postsynaptic targets, suggesting that FRMPD2B has contributed to the expansion of cortical connectivity during human evolution.

**Fig. 2.**
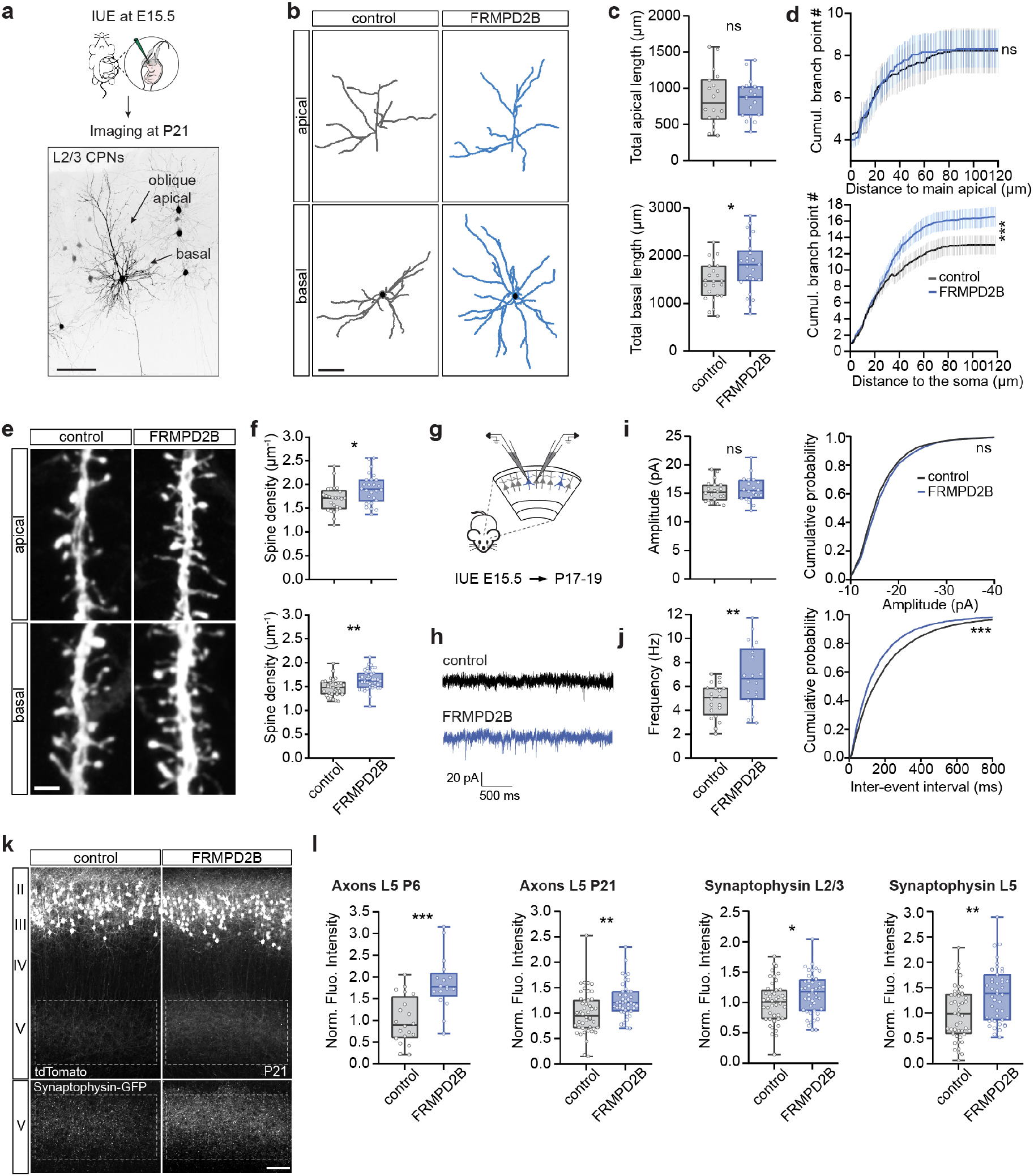
FRMPD2B regulates L2/3 CPN morphological complexity and connectivity. **a**, Experimental design: Cortical progenitors generating L2/3 CPNs were transfected by in utero electroporation (IUE) at embryonic day 15.5 (E15.5). Their apical and basal dendrites were imaged at postnatal day 21 (P21) in brain slices using confocal microscopy. Scale bar: 50 µm. **b**, Representative reconstructions of apical and basal dendritic arbors. Scale bar: 50 µm. **c**, Total dendritic length of apical (top) and basal (bottom) dendrites. Apical: n_control_=17, n_FRMPD2B_=19; basal: n_control_=23, n_FRMPD2B_=25 neurons. **d**, Cumulative branchpoint number in the indicated conditions. Error bars represent SEM. **e**, Confocal images of dendritic spines in basal and apical dendrites in control and FRMPD2B-expressing neurons. Scale bar: 2 µm. **f**, Quantification of spine density in apical and basal dendrites. Apical: n_control_=25, n_FRMPD2B_=26; basal: n_control_=32, n_FRMPD2B_=39 neurons. **g**, Whole-cell patch clamp recording in acute brain slices at P17-19 following IUE at E15.5. **h**, Representative mEPSC traces of control and FRMPD2B-expressing neurons. **i**, Mean mEPSC amplitude and cumulative distribution of the amplitude of the first 250 events per cell, n_control_=20, n_FRMPD2B_=20. **j**, Mean mEPSC frequency and cumulative distribution frequency of the inter-event interval of the first 250 events per cell. **k**, Coronal brain section of P21 mouse cortex electroporated at E15.5 with plasmids encoding tdTomato (top) and synaptophysin-GFP (bottom) with or without FRMPD2B. Numbers on the left indicate cortical layers. Dashed lines indicate the quantified area. L6 was not depicted in the images. Scale bar: 100 µm. **l**, Quantification of axonal branching (top) and presynaptic bouton density (bottom) according to tdTomato and synaptophysin-GFP fluorescence intensity in the indicated conditions. P6: n_control_=19 slices; n_FRMPD2B_=17 slices (6 mice per condition); P21: n_control_=46 slices (11 mice); n_FRMPD2B_=45 slices (10 mice). NS,*P>0*.*05*, ^*^*P<0*.*05*, ^**^*P<0*.*01* and ^***^*P<0*.*001* using Mann-Whitney test (c, f, l), two-way repeated measures ANOVA (d), Student t-test (i, j, l) and Kolmogorov-Smirnov (cumulative distributions in d, i, j).

### Regulation of FRMPD2B by activity and experience

Human CPNs are characterized by their extremely slow tempo of morphogenesis (their neoteny) compared to that of other species^3^. The increase in dendritic complexity and synapse number induced by *FRMPD2B* suggested that this gene is only transcribed when neurons have reached a certain level of activity during maturation. To test for activity-dependent regulation, we designed a reporter assay in which EGFP expression was driven by the 1 kb sequence upstream of the transcription start site (TSS) of *FRMPD2B* (referred to as pFRMPD2B) (Fig. 3a), which is expected to contain the promoter region. We introduced this reporter in mouse upper layer CPNs using IUE and quantified the fluorescence associated with EGFP normalized to the level of cytosolic tdTomato expressed under a ubiquitous, chicken beta-actin promoter. pFRMPD2B induced EGFP expression and its transcriptional activity increased mildly between P1 and P7, then sharply between P7 and P14 (Fig. 3b), paralleling the trajectory of synaptogenesis. We then assessed if pFRMPD2B is regulated by experience using a paradigm of sensory deprivation consisting of daily bilateral whisker trimming (Fig. 3c). At P14, the activity of pFRMPD2B in upper layer CPNs of the somatosensory cortex was lower in sensory-deprived mice than in their control littermates (Fig. 3c, d). Conversely, increasing sensory, social and cognitive experiences with environmental enrichment (EE) from E10 to P14 strongly increased pFRMPD2B-driven EGFP expression compared to standard housing conditions (Fig. 3c, d and Fig. S5). In line with this, EGFP-associated fluorescence intensity in L2/3 CPNs was decreased by overexpression of the potassium channel Kir2.1, and markedly increased by overexpression of the sodium channel mNaChBac (Fig. 3e, f). Thus, the transcriptional activity of *FRMPD2B* promoter is bidirectionally regulated by experience-dependent changes in neuronal activity.

**Fig. 3.**
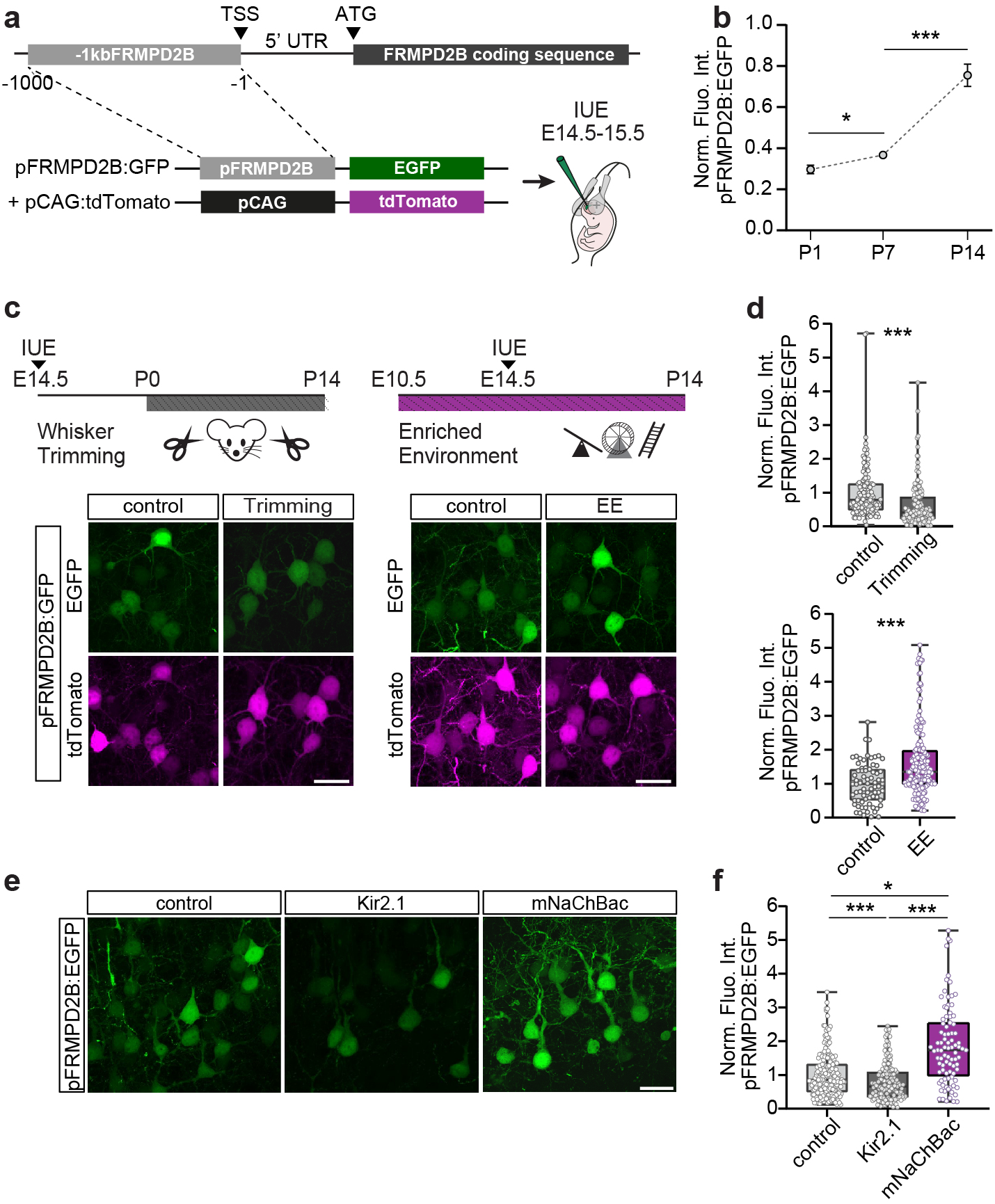
Bi-directional regulation of *FRMPD2B* promoter activity by experience and neuronal activity. **a**, Experimental design: *FRMPD2B* promoter activity was evaluated in upper-layer CPNs targeted by IUE with a reporter assay in which EGFP expression was driven by the 1-kb genomic sequence upstream of the transcription start site (TSS). The expression level of tdTomato driven by a CAG promoter was used for normalization. **b**, Normalized fluorescence intensity of EGFP under pFRMPD2B at different developmental timepoints. Error bars represent SEM. n_P1_=70 cells (4 mice), n_P7_=161 cells (5 mice), n_P14_=128 cells (6 mice). **c**, Representative confocal images of upper layer CPNs expressing the reporter constructs at P14 in control condition, after whisker trimming and in enriched environment (EE) conditions. Scale bar: 20 µm. **d**, Normalized fluorescence intensity of EGFP in the indicated conditions. Trimming: n_control_=128 cells (6 mice), n_trimming_=141 cells (8 mice), EE: n_control_=76 cells (5 mice), n_EE_=176 cells (10 mice). **e**, Confocal images of L2/3 CPNs expressing the reporter in control condition or together with Kir2.1 or mNacChBac at P7. Scale bar: 10 µm. **f**, Quantification of EGFP normalized fluorescence intensity in the indicated conditions. n_control_=187 cells (9 mice), n_Kir2.1_=145 cells (9 mice), n_mNaChBac_=87 cells (5 mice). ^*^*P<0*.*05*, ^***^*P<0*.*001* using Kruskal-Wallis with Dunn’s multiple comparisons test.

### Gain of activity-dependent regulation through a retrotransposon insertion

The promoter region of *FRMPD2B* resides within the duplication, raising the question of how its regulation differs from that of its ancestral copy. We tested whether the activity-dependent regulation is a unique feature of *FRMPD2B* promoter by assessing, with the reporter assay, how sensory deprivation impacts the putative transcriptional activity of the ancestral paralogous sequence (phumPara) and macaque orthologous sequence (pmacOrtho) (Fig. 4a). Both sequences were able to drive EGFP expression but neither of them was sensitive to experience (Fig. 4b, c). Therefore, the emergence of activity-dependent responsiveness in *FRMPD2B* promoter postdates the duplication. Using sequence analysis, we found that although pFRMPD2B and phumPara are highly identical (99.6%), pFRMPD2B contains a distinct 303 bp insertion located 124 bp upstream of the TSS (Fig. 4d). This insertion corresponds to an Alu repeat of the AluYe5 subfamily. Alu repeats are primate-specific retrotransposons that account for approximately 10% of the modern human genome and have been implicated in the evolution of gene regulation^43–45^. AluYe5 retrotransposons belong to one of the most recent subfamilies and are only found in great apes^46^. Using *in silico* transcription factor binding site (TFBS) analysis and position weight matrices, we found that the AluYe5 insertion in pFRMPD2B contains binding motifs for the activity-regulated transcription factors MEF2, FEZF2 and KLF10 (Fig. 4d and Table S1). These TFBS were unique to the retrotransposon insertion and were not detected elsewhere in the *FRMPD2B* promoter region, in the paralogous ancestral sequence, or in the orthologous macaque sequence. Deletion of the MEF2 binding site in pFRMPD2B abolished both the developmental increase in promoter activity and its experience-dependent regulation (Fig. 4e). Thus, *FRMPD2B* evolved in two steps: first the ancestral gene was partially duplicated in the human lineage, then a retrotransposon insertion modified the promoter architecture of the human-specific copy and introduced a MEF2 binding site that conferred activity- and experience-dependent regulation to the human-specific gene (Fig. 4f).

**Fig. 4.**
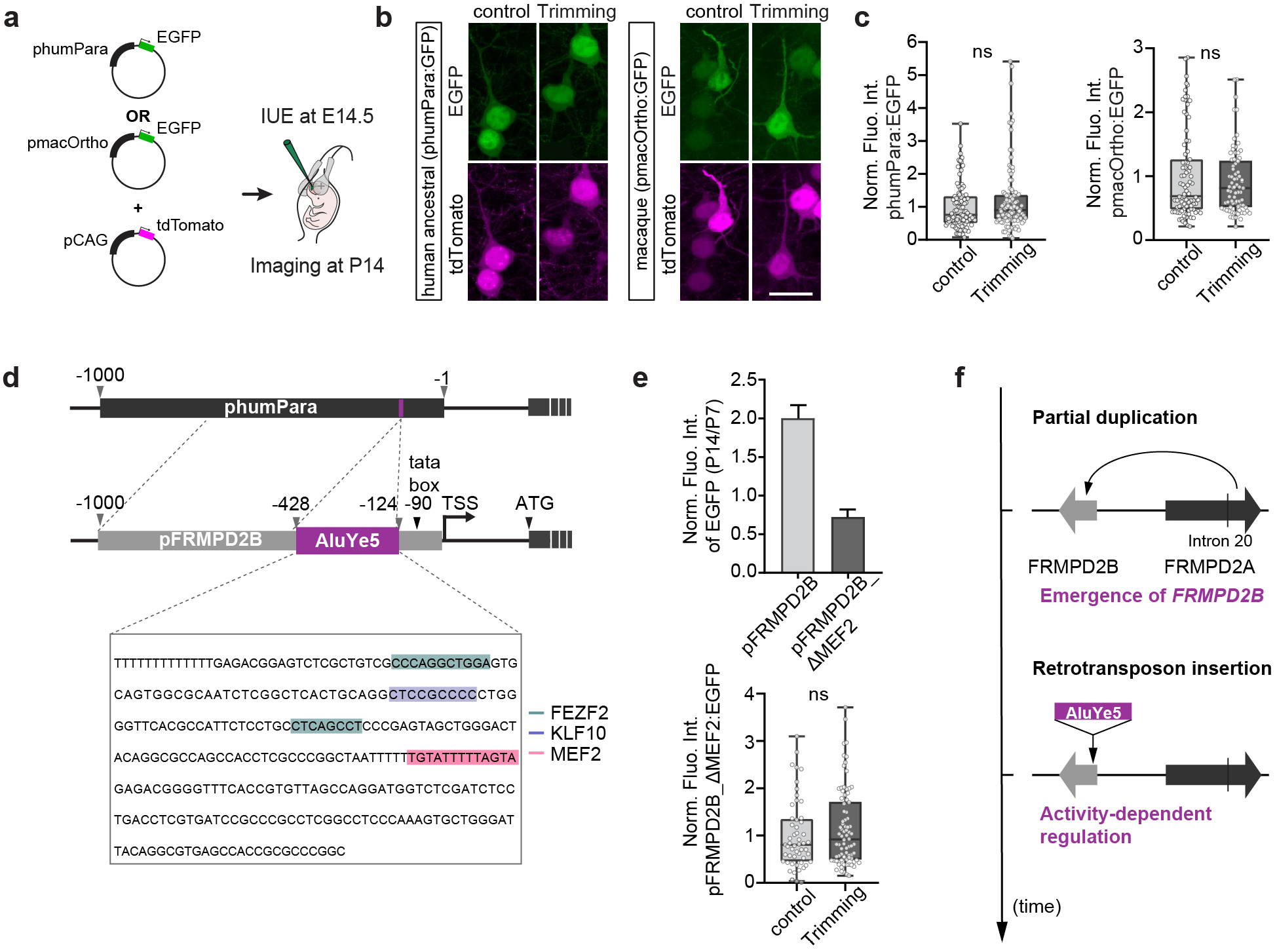
Evolutionary history of *FRMPD2B* experience-dependent regulation. **a**, Experimental design: the reporter assay was used to test the experience-dependent regulation of the ancestral paralogous sequence (phumPara) and macaque orthologous sequence (pmacOrtho) of pFRMPD2B **b**, Confocal images of L2/3 CPNs expressing the reporter constructs in control and after whisker trimming. Scale bar: 20 µm. **c**, Normalized EGFP fluorescence intensity in the indicated conditions, phumPara: n_control_=98 cells, n_trimming_=108 cells (7 mice per condition); pmacOrtho: n_control_=87 cells, n_trimming_=77 cells (5 mice per condition). **d**, Alignment of phumPara and pFRMPD2B. Upstream of the TATA box, pFRMPD2B contains a unique insertion corresponding to an AluYe5 retrotransposon providing binding sites for activity-dependent transcription factors. TSS: transcription start site. **e**, Impact of MEF2 binding site deletion (ΔMEF2) on the developmental and experience-dependent regulation of pFRMPD2B activity. Developmental regulation between P7 and P14: pFRMPD2B: n_P7_=161 cells, n_P14_=128 cells (5 and 6 mice); pFRMPD2B_ΔMEF2: n_P7_=23 cells, n_P14_=48 cells (2 and 4 mice). Whisker trimming: n_control_=53 cells, n_trimming_=82 cells (4 and 5 mice). Error bars represent SEM. **f**, Schematic of the evolutionary history of *FRMPD2B*. The partial duplication of the ancestral gene *FRMPD2A* in the human lineage gave rise to the human-specific gene *FRMPD2B*. Subsequently, the insertion of an AluYe5 retrotransposon in the promoter region of *FRMPD2B* conferred activity-dependent regulation. NS,*P>0*.*05* by Mann-Whitney test.

### FRMPD2B is required for human CPN morphological complexity

We next assessed the role of FRMPD2B directly in human CPNs. To do so, we used a model of xenotransplantation in which CPNs derived from human pluripotent stem cells (hPSCs) integrate into the mouse cortex. The xenotransplantation promotes the morphological, transcriptomic and electrophysiological maturation of human CPNs, while preserving their species-specific developmental tempo^47–50^. The xenotransplantation also enables these neurons to express activity-regulated genes critical for cortical circuit maturation, such as *BDNF* and *NPTX2*^50^, or *FRMPD2B* (Fig. S6). By contrast, when maintained *in vitro*, hPSC-derived CPNs typically fail to progress beyond the mid-fetal stage and poorly express activity-regulated genes^50,51^. Cortical organoids were cultured for 8 to 10 weeks in vitro^52^. Then, cells were dissociated, plated and cultured in 2D. After 2 days in vitro, cells were transduced with lentiviruses expressing an shRNA against *FRMPD2B* or a control shRNA, and EGFP for identification and morphological analysis (Fig. 5a and Fig. S7e). On the next day, they were treated with a g-secretase inhibitor (DAPT) and antimitotic drug (Cytarabine, ARA-C) to promote neurogenesis and eliminate progenitors^48^. This protocol enabled the enrichment of upper layer CPNs expressing CUX1/2 (Fig. S7a, b), in which *FRMPD2B* is primarily expressed. The human cells were transplanted into the host mouse brain in utero at E14.5-E15.5, when mouse upper layer CPNs are generated and the structures necessary to support neuronal migration are in place. One month post-transplantation (1 MPT), human CPNs mostly populated the upper layers of the mouse cortex (Fig. 5b) and expressed upper-layer CPN identity marker (Fig. S7c). Despite similar distance from the pial surface, as measured by the length of the main apical dendrite, human CPNs knocked down for *FRMPD2B* with two distinct shRNAs had a much simpler morphology, with less branched and shorter dendrites than control neurons (Fig. 5c, d and Fig. S7e). *MEF2* knockdown had similar effects on dendritic length and branching (Fig. 5c, d and Fig. S7e), suggesting that FRMPD2B is a human-specific effector of MEF2. These data demonstrate that FRMPD2B is necessary for the proper morphogenesis of human upper layer CPNs.

**Fig. 5.**
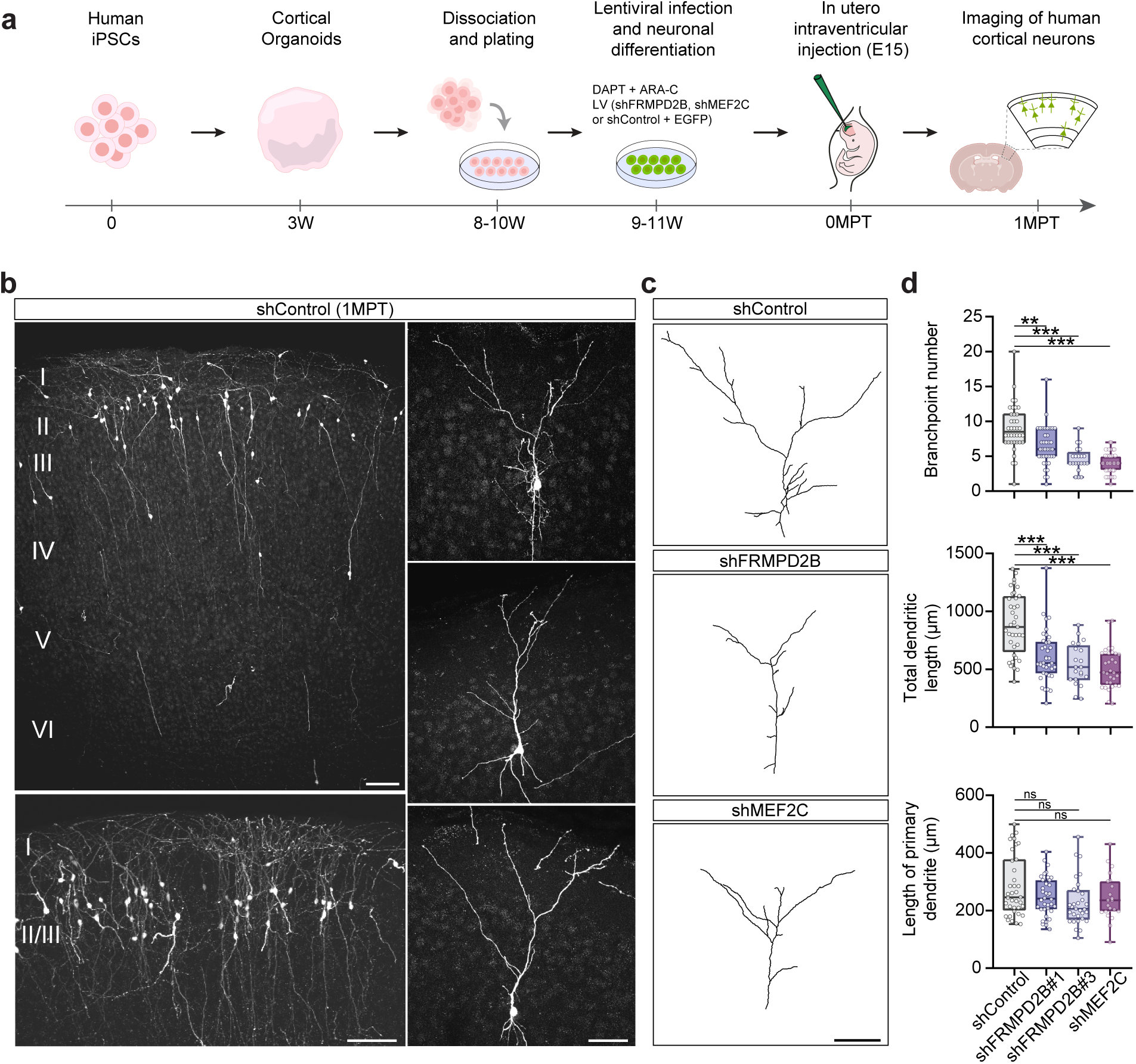
FRMPD2B is required for complex dendritic arbor development in xenotransplanted human CPNs. **a**, Experimental design: Cortical organoids were generated from human induced pluripotent stem cells (hiPSCs). After 8-10 weeks of differentiation, organoids were dissociated, plated and infected with lentiviral vectors expressing EGFP together with either shControl, shFRMPD2B, or shMEF2C. DAPT and ARA-C were added to the culture medium in order to promote neurogenesis and eliminate proliferating progenitors. Human CPNs were then transplanted as single cells into the lateral ventricles of E15 mouse embryos by in utero intraventricular injection. EGFP-positive xenotransplanted human CPNs were imaged and analyzed 1 month post-transplantation (1 MPT). **b**, Left: representative images of human CPN distribution across the mouse somatosensory cortex showing preferential localization to L2/3 (scale bars: 100 µm; lower inset: zoom on L2/3). Right: representative morphology of xenotransplanted human CPNs at 1 MPT (scale bar: 50 μm). **c**, Representative reconstructions of apical dendritic arbors in shControl, shFRMPD2B, or shMEF2C conditions. Scale bar: 50 µm. **d**, Quantification of branchpoint number, primary and total dendritic length of apical dendrites. n_shControl_=37 cells (9 mice), n_shFRMPD2B#1_=38 (9 mice), n_shFRMPD2B#3_=22 cells (4 mice), n_shMEF2C_=31 (4 mice). NS,*P>0*.*05*, ^**^*P<0*.*01*, ^***^*P<0*.*001* using Kruskal-Wallis with Dunn’s multiple comparisons test and One-way ANOVA with Tukey’s multiple comparisons test.

## DISCUSSION

Our results identify *FRMPD2B* as a human-specific and experience-regulated gene whose expression peaks during the first years of life and is restricted to cell types mediating cortico-cortical connectivity. We show that FRMPD2B expression in mouse upper layer CPNs promotes dendritic and axonal development, increases dendritic spine and presynaptic bouton density and enhances functional synaptic connectivity, thereby contributing to distinct features of human CPNs. We find that *FRMPD2B* promoter activity is bidirectionally regulated by sensory deprivation and environmental enrichment, as well as by neuronal activity. This activity-dependent regulation is unique to the human-specific gene: neither the paralogous ancestral sequence of the promoter region nor its orthologous macaque counterpart is sensitive to experience. *FRMPD2B* acquired responsiveness to experience after the duplication through which it was generated, when the insertion of an AluYe5 retrotransposon into its promoter introduced a binding site for MEF2 transcription factors. Xenotransplantion of human upper layer cortical pyramidal neurons into the mouse brain established the critical role of FRMPD2B in human neuronal morphogenesis and strengthened its link to MEF2, a major ID/ASD risk gene. Our results reveal how successive human-specific genetic innovations have endowed human cortical neurons with species-specific response to experience enabling the elaboration of their complex morphology and the expansion of their connectivity during postnatal development.

Despite growing understanding of human uniqueness at the genomic level, the contribution of human-specific genetic innovations to postnatal cortical development and their potential interactions with activity-regulated mechanisms have remained largely inaccessible to experimental investigation. Pediatric human brain samples, particularly in the first years of life, are scarce for obvious reasons, while human cortical neurons derived from hPSCs require xenotransplantation into the rodent brain to reach a compatible stage of maturation and recapitulate the expression of activity-regulated transcriptional programs^3^. To address this experimental challenge and determine FRMPD2B function in human neurons, we developed a xenotranspnatation approach enabling the in vivo maturation and genetic manipulation of human upper layer cortical pyramidal neurons, in which FRMPD2B is predominantly expressed. Because of the slow tempo of human corticogenesis, previous xenotransplantation approaches had so far mostly relied on early-born, deep-layer neurons^47–49,53^. Our approach now provides experimental access to late-born, upper layer CPNs, which have extensively expanded and diversified during human evolution^54,55^. Using this system and building on the results obtained in mice, we directly demonstrate that FRMPD2B is necessary for human CPNs to develop their complex morphology. Previous studies have shown that the human-specific genes SRGAP2B/C delay synaptic maturation and thereby extend the postnatal period of synaptic development^32,48,53,56^. Our work now highlights that *FRMPD2B* endows human neurons with human-specific mechanisms that shape their response to postnatal experience. While the expansion of cortico-cortical connectivity in humans has so far been viewed primarily through the lens of neurogenesis and intrinsic mechanisms^32,33,57–59^, our results suggest a distinct but complementary evolutionary strategy through which postnatal experiences drives the increase in connectivity.

It was previously suggested that FRMPD2B might operate by inhibiting its ancestral copy FRMPD2A^60^, a protein necessary for the maintenance of polarity in epithelial cells^61^ and involved in synapse specification in the retina^62^. However, both our long-read RNA sequencing and public RNAseq databases indicate that FRMPD2B and long ancestral FRMPD2A are not co-expressed in the human neocortex. Although we cannot rule out that short transcripts originating from the ancestral gene, indistinguishable from *FRMPD2B* transcripts, are also expressed in the neocortex, the developmental and experience-dependent regulation is unique to *FRMPD2B*. Thus, *FRMPD2* expression profile in the human neocortex is shaped by the evolutionary history of *FRMPD2B*, from the initial duplication to the retrotransposon insertion through which it gained activity-responsiveness. The developmental and experience-dependent increases in *FRMPD2B* expression in cortical neurons require the activity-regulated transcription factors MEF2. Consistently, we found that both *MEF2C* and *FRMPD2B* inactivation decreased dendritic length and branching in xenotransplanted human CPNs. Studies in mice have shown that MEF2 transcription factors regulate neuronal survival, axonal and dendritic development, and cortical circuit connectivity during postnatal development and critical periods of plasticity^63,64^. In the human neocortex, *MEF2A* exhibits a neotenic expression pattern compared to macaques and chimpanzees, whereas *MEF2C* is associated with a human accelerated region (HAR), both of which have been linked to ASD^65,66^. Together with evidence of recurrent deletions and duplications associated with neurodevelopmental disorders in the region encompassing *FRMPD2* duplication^34^, this posits the FRMPD2B-MEF2 axis as critical for human brain development and evolution. Remarkably, the human-specific binding site for MEF2 in *FRMPD2B* promoter is provided by a primate-specific AlueYe5 retrotransposon inserted upstream of the TSS. AlueYe5 elements exist in thousands of copies in the human genome. They may therefore confer primate- or human-specific activity-dependent regulation to other genes and further impact brain development and plasticity^67^.

The discovery of *FRMPD2B* as an activity-regulated human-specific gene links the evolution of the human genome to the plasticity of the human brain and highlights how, at the cellular and molecular levels, the development of the human neocortex became increasingly dependent on environmental interactions during evolution. It also raises new questions on the pathophysiology of neurodevelopmental disorders featuring early excitation/inhibition imbalance, including some forms of ASD, ID and epilepsy^23,68^, in which *FRMPD2B* could influence the phenotypic outcome in a way that cannot be modelled in mice or in human cells in vitro. Determining whether and how human-specific modifiers of neuronal and synaptic development impact these disorders, and deciphering the human-specific aspects of their pathophysiology will be critical for future therapeutic strategies.

## METHODS

### Animals

All animals were handled according to French and EU regulations (directive 2010/63, decree 2013-118). Authorization references were #1530-2015082611508691v3, #29476 2021020311595454v5, and #60645-2026041511178520v10 delivered by the French Ministry of Research after evaluation by the Comité d’Ethique en Experimentation Animale n°005. In utero electroporation experiments were conducted on pregnant Rj:SWISS females at E14.5-15.5 (Janvier labs) and offsprings were used for experiments at indicated timepoints without consideration of their sex. Mice were maintained in a 12 hours light/dark cycle with unlimited access to food and water.

### DNA constructs

The human FRMPD2B cDNA (BC071635, clone #5243609 from Dharmacon) was subcloned between XhoI and KpnI into pCAG with C-terminal tags of EGFP or 3XHA (vector described before^69^. pCAG mVenus and pCAG tdTomato were previously described^69^. pCAG Flex mVenus was obtained by PCR amplification of mVenus in pCAG mVenus, then subcloned into AAV pCAG Flex-tdTomato WPRE (addgene # 51503, gift from Hongkui Zeng) between the BsrGI and XhoI restriction sites. pCAG Synaptophysin-GFP was obtained by PCR amplification of Synaptophysin sequence from pAAV FLEX Synaptophysin GFP (addgene #137188, gift from Matthew Nolan) then inserted between XhoI and KpnI restriction sites into pCAG with a C-terminal tag of EGFP. pCAG Cre was obtained from X. Morin (IBENS, Paris, France). pCAG Kir2.1-T2A-tdTomato and pCAG mNaChBac-T2A-tdTomato were a gift from Massimo Scanziani (Addgene plasmid #60598; RRID: Addgene_60598; Addgene plasmid #60650; RRID: Addgene_60650). The 1kb sequences upstream of the transcription start sites of *FRMPD2B* (T2T-CHM13v2.0 chr10: 47.764.423 - 47.765.423) and putative short transcripts of human and macaque *FRMPD2A* (T2T-CHM13v2.0 chr10: 49.024.136 - 49.025.240 and Mmul_10 - rheMac10 chr9: 49.202.244 -49.203.244, respectively) were produced through gene synthesis by Integrated DNA technologies (IDT). The mutant pFRMPD2B sequence in which the MEF2 binding site was deleted was produced through gene synthesis by Eurofins genomics. All putative promoter constructs were cloned and inserted in front of EGFP between MIuI and AgeI into a pCAG vector by replacing the chicken β-actin promoter. For knockdown with lentivirus, we used a lentiviral vector (pLV) expressing shRNAs under an H1 promoter and EGFP under a synapsin promoter^69^. The seed sequences were 5’- ACTACCGTTGTTATAGGTG-3’ for shControl, 5’-CCAGACTTTCGACAGATATTT-3’ for shFRMPD2B and 5’- GCTGTGTGACTGTGAGATTGC-3’ for shMEF2C inserted at the XhoI/XbaI sites. All constructs were verified by DNA sequencing.

### Lentivirus production

HEK293T cells were co-transfected with the lentiviral vector expressing the shRNAs and two plasmids together comprising the lentiviral packaging system. For virus concentration, the viral supernatant was collected 48 h after transfection and ultracentrifuged at 4°C at 20,000 g for 2 h. Viral pellets were resuspended in sterile 0.1% BSA in PBS, aliquoted and stored at -80°C. Titer check was performed for every batch. 100,000 cells were seeded on gelatin-coated coverslips (12-well plates) and infected with 6 serial dilutions of the lentivirus. Medium was changed 24 h later, and cells were fixed 72 h later for 1 h at room temperature with PBS PFA 4%, followed by immunostaining of the relevant marker and Hoechst. From this, the volume of virus needed to infect a given number of cells was estimated.

### shRNA validation

shRNAs were validated in mycoplasma-negative HEK293T cells (ATCC Cat# CRL-1573; RRID: CVCL_0045). Briefly, HEK293T cells were co-transfected using Jet-Prime (Polyplus Transfection #POL114-07) according to the manufacturer protocol with either HA-tagged FRMPD2B or HA-tagged MEF2C together with the shRNAs targeting them or a control shRNA at a 1:3 ratio. Two days after transfection, cells were collected and lysed in RIPA buffer (150 mM NaCl, 1.0% NP-40, 0.5% sodium deoxycholate, 0.1% SDS, 50 mM Tris, pH 8.0, Sigma-Aldrich) and further processed for western blot analysis of the relative protein expression levels.

### In utero electroporation and slice preparation

In utero electroporation (IUE) was performed as previously described^32,69^. Pregnant Swiss females at E14.5-15.5 (Janvier Labs) were injected subcutaneously with 0.1 mg.kg^-1^ buprenorphine for analgesia and anaesthetized with isoflurane (4% for induction and 2-2.5% during surgery; ISO-VET Piramal Critical Care). The uterine horns were exposed after laparotomy to allow injection of the endotoxin-free DNA mixture into a ventricle of mouse embryos. The injected embryos were then electroporated with 4 pulses of 40 V for 50 ms at 500 ms intervals generated by a square wave electroporator (Sonidel #CUY21) and delivered by a platinum disc electrode forceps (5 mm diameter, Sonidel # CUY650P5). Subsequently, the uterus was placed back into the abdominal cavity and the incision was sutured. Plasmids were diluted in 1X PBS (Sigma #D8537) and 0.1% Fast Green (Sigma #F7252) dye to allow visualization. The volume of the injected DNA mixture was adjusted according to the experiments. Animals were sacrificed at the indicated age by intracardiac infusion of 4% paraformaldehyde (Electron Microscopy Sciences # 15714-S) in PBS. Unless otherwise indicated, 100 μm coronal brain sections were obtained using a vibrating microtome (Leica VT1200S, Leica Micro-systems). Sections were mounted on slides in Vectashield medium (Eurobio #H-1000).

### Ex utero electroporation and primary neuronal culture

Ex utero electroporation was performed as previously described^32^ using timed pregnant Rj:SWISS female mice at E15.5. After injection of endotoxin-free DNA with a glass pipette into lateral ventricles, embryonic mouse brains were electroporated using a square wave electroporator (ECM 830, BTX) and tweezer-type platinum disc electrodes (5 mm-diameter, Sonidel # CUY650P5). The electroporation settings were: 5 pulses of 20 V for 100 ms with 200 ms intervals. After electroporation cortices were dissected and dissociated using papain (Worthington, LK003178). Neurons were plated on poly-L-ornithine (80 μg/ml, Sigma) in Neurobasal™ (Gibco #21103049) supplemented with GlutaMAX™, B27 (Gibco #17504044) and penicillin (2.5 units/ml)-streptomycin (2.5 μg/ml) (Gibco#15070063). Cells were maintained at 37°C in 5% CO2 until use.

### Generation of cortical organoids

Cortical organoids were generated from mycoplasma-negative human induced pluripotent stem cells (iPSCs) (HMGU#1, ISFi001-A, RRID:CVCL_YT30) as previously described^52^. Day 0 to Day 4: iPSC colonies were detached using pre-warmed collagenase (1 mg/ml) for 45 minutes at 37°C. The floating colonies were transferred using a cut pipette tip into a 15 ml tube and washed with Medium 1 (DMEM-F12 without phenol red, 20% KnockOut Serum Replacement (KSR), 1× GlutaMAX, 1× MEM Non-Essential Amino Acids (MEM-NEAA), 1× 2-mercaptoethanol, penicillin/streptomycin, 2 μM Dorsomorphin, 2 μM A-83). The colonies were then transferred into ultra-low attachment plates with 3 ml of Medium 1 and cultured at 37°C in a 5% CO2 atmosphere. Day 5 to Day 6: Half of the Medium 1 was replaced daily with Medium 2 (DMEM-F12 without phenol red, 1× N2 supplement, 1× GlutaMAX, 1× MEM-NEAA, penicillin/streptomycin, 1 μM CHIR-99021, 1 μM SB-431542). Day 7 to Day 14: On Day 7, embryoid bodies (EBs) were embedded in Matrigel diluted in Medium 2 at a ratio of 2:1. The Matrigel-EB mixture was spread into regular 6-well plates and incubated at 37°C for 30 minutes to solidify (10–20 EBs per well). Medium 2 was gently added to the well without disturbing the Matrigel patch. On Day 14, the Matrigel-EB mixture was mechanically dissociated using a 5 ml pipette and transferred into a 15 ml tube for gentle washing. Organoids were then suspended in Medium 3 (DMEM-F12 without phenol red, 1× N2 supplement, 1× B27 supplement (with vitamin A), 1×GlutaMAX, 1×MEM-NEAA, 1× 2-mercaptoethanol, penicillin/streptomycin, 2.5 μg/ml insulin) and cultured in ultra-low Attachment 6-well plates under agitation at 100 rpm using a Digital Orbital Shaker (DOS-10M from ELMI). Day 35 to Day 84: Starting from Day 35, Medium 3 was supplemented every two days with diluted Matrigel (1:100) to support the growth and development of the cortical organoids.

### In utero transplantation of human cortical neurons in the mouse cortex

Human cortical organoids of 8-10 weeks of maturation were dissociated using a Neurosphere Dissociation Kit (Miltenyi Biotec, #130-095-943) and plated in Matrigel in Neural Stem Cell (NSC) Medium composed of DMEM-F12 without phenol red (Gibco, #21041025) containing D-glucose (2.9 mg.ml^-1^), 1xB27 supplement without vitamin A (Gibco, #12587010), penicillin/streptomycin 5 U.ml^-1^, amphotericin B 250 ng.ml^-1^, 20 ng.ml^-1^ fibroblast growth factor (FGF), 20 ng.ml^-1^ epidermal growth factor (EGF). On DIV1-2, cells were infected with the lentiviral vector. After 24 hours, cells were washed with NSC medium twice and were treated with 10 µM DAPT (Abcam, Cat#ab120633) for 72 hours, and then 10 µM DAPT and 5 µM Cytarabine (ARA-C) (Merck, Cat#C3350000) for 24 hours. The day of the transplantation, cells were dissociated using Accutase (StemCell Technologies, #07920) and suspended in the injection solution containing 20 mM EGTA (Merck, Cat#03777) and 0.1% Fast Green (Merck, Cat#210-M) in PBS at 250,000 cells.ml^-1^. Laparotomy procedure of Swiss pregnant mice at E15 was carried out as described above for in utero electroporation and transplantation was achieved through the injection of approximately 1 μl of cell suspension into the lateral ventricles of the embryos. Data were obtained from 3 (shFRMPD2B#1), 1 (shFRMPD2B#3) and 2 (shMEF2C) independent batches of organoids.

### Immunostaining of human cortical neurons

Newly generated neurons cultured on coverslips were fixed in 4% paraformaldehyde (PFA) in PBS for 15 min at RT. Cells were then permeabilized with 0.1% Triton X-100 in PBS for 4 min and blocked in 10% bovine serum albumin (BSA) in PBS for 30 min at room temperature. Samples were incubated with primary antibodies diluted in blocking solution overnight at 4°C. Following three washes with PBS, cells were incubated with the appropriate secondary antibodies for 45 min at RT. Coverslips were subsequently washed three times with PBS prior to mounting and imaging. Xenotransplanted animals were transcardially perfused with 4% PFA in PBS. Brains were dissected and post-fixed overnight at 4°C in the same fixative solution. Following post-fixation coronal brain sections (100 μm thick) were obtained. For immunofluorescence staining, free-floating sections were incubated in blocking solution consisting of 0.3% Triton X-100, 5% horse serum (HS), and 3% BSA in PBS for 2 h at room temperature. Sections were then incubated with primary antibodies diluted in blocking solution overnight at 4°C with gentle agitation. Following three washes in PBS, sections were incubated for 2 h with the secondary antibodies diluted in blocking solution PBS. Finally, sections were washed three times in PBS prior to mounting and imaging. The following primary antibodies were used: chicken anti-EGFP (1:2000; AB_2307313, Aves Labs), rat anti-BCL11B/CTIP2 (1:500, ab18465, Abcam), rabbit anti-Cux1/2 (1:500, ab309139, Abcam), rabbit anti-NeuN (1:500, ab177487, Abcam), mouse anti-HNA (1:200, ab191181, Abcam), goat anti-SOX2 (1:500; AF2018, R&D Systems,) mouse anti-βIII-tubulin (TUBB3; TUJ1; 1:1000; 801201, BioLegend).

### Environmental enrichment and whisker trimming

The environmental enrichment (EE) paradigm was adapted from previously established protocols. ^70,71^. Pregnant Swiss mice (3 for enrichment, 1 for control) were randomly assigned to control (37 x 23 x 14 cm) and EE (Marlau™ Viewpoint, Civrieux, 80 × 60 × 51 cm) cages at E10.5. Following in utero electroporation and delivery, pups remained in their respective cages until P14. In addition to the standard Marlau cage configuration, enriched housing contained objects of different shapes, colours and textures that were replaced daily and all together once per week. Nesting material (paper houses and shredded paper) was also provided. In control, mice were kept in standard housing conditions and were maintained at a maximum of 6 pups in each cage. Whisker trimming was performed from P0 to P14 for half of the mice in each litter. All whiskers were trimmed bilaterally every day to approximately 1 mm from the skin surface. Trimming was carried out by gently restraining the pups in the experimenter’s hand. To ensure consistency, pups in the control group underwent the same gentle handling and restraint procedure each day, but their whiskers were left intact.

### Slice preparation for electrophysiology

Following in utero electroporation, swiss juvenile mice (postnatal day 9-12 or 17-19) were deeply anesthetized with isoflurane (ISO-VET, Piramal Critical Care) and brains were quickly removed and placed in cold (4 °C), oxygenated (95% O2 and 5% CO2) cutting solution, containing: 85 mM NaCl, 64 mM sucrose, 25 mM glucose, 2.5 mM KCl, 1.2 mM NaH_2_PO_4_, 24 mM NaHCO_3_, 0.5 mM CaCl_2_, and 7 mM MgCl_2_ (300-310 mOsm, pH 7.3–7.4). Acute coronal slices (300 μm thickness) were sectioned using the 7000 smz-2 vibratome (Campden Instruments) and left to recover for approximately 45 minutes at room temperature in oxygenated artificial cerebrospinal fluid (ACSF) containing: 125 mM NaCl, 2.5 mM KCl, 2 mM CaCl_2_, 1 mM MgCl_2_, 1.2 mM NaH_2_PO_4_, 24 mM NaHCO_3_, and 25 mM glucose (300-310 mOsm, pH 7.4). Slice recordings were performed in a submerged recording chamber at 34°C, with continuous perfusion with oxygenated ACSF at a rate of 4 ml/min.

### Electrophysiological recordings

Whole-cell voltage- and current-clamp recordings were acquired from visually identified electroporated or non-electroporated layer 2/3 cortical pyramidal neurons of the somatosensory cortex. Slices were visualized using an IR-DIC microscope (Olympus), equipped with a 4X objective (UPlanFL, NA 0.13) as well as a 40X water immersion objective (LUMPlan Fl/IR, NA 0.8) and a ORCAFlash 4.0LT camera (Hamamatsu). Signals were recorded with a Multiclamp 700B Amplifier (Axon instruments), Axon Digidata 1550 (Axon Instruments) and pClamp 10.7 software (Axon instruments). Intrinsic excitability and mEPSCs were recorded through borosilicate glass pipettes (3-6 MΩ) filled with an intracellular solution, containing: 144 mM K-gluconate, 7 mM KCl, 10 mM HEPES, 1 mM EGTA, 1.5 mM MgCl_2_, 2 mM NaATP, 0.5 mM NaGTP, (295 mOsm, pH 7.3 adjusted by KOH). To isolate mEPSCs, cells were voltage-clamped at -60 mV while ACSF was supplemented with 0.5 µM TTX (Tocris, #1069) and 10 µM gabazine (Abcam, #ab120042). All data were sampled at 10 kHz and filtered at 2 kHz. Traces were analyzed offline over 1-minute periods using pClamp 10.7 (Molecular Devices). Cells showing >20% change in access and input resistance upon application of 5 mV hyperpolarizing steps of current were excluded from the analysis.

### Confocal image acquisition and analysis

Confocal images were acquired using a Nikon A1R HD25 scanning confocal microscope, controlled by NIS Elements and equipped with an LU-N4/N4S 4-laser unit that includes 405 nm, 488 nm, 561 nm, and 640 nm lasers. The objective lenses used were the Plan Apo Lambda 10x Air NA 0.45 (MRD00105) for the identification of electroporated neurons and whole neuron images, the CFI Plan Apochromat Lambda D 60X Oil and the SR HP Plan Apo Lambda S 100x Silicone NA 1.35 (MRD73950) (Nikon) for high magnification images. Z-stacks were acquired with 150 nm steps for spine morphology and localization, and a 1 µm spacing for dendrite morphology and axon quantification. The acquisition parameters for dendritic spine analysis were set to 1024X1024 mode, a pixel size of 100 nm, an exposure rate of 2.4 per pixel, and images were averaged over 2 acquisitions. Parameters for whole neuron reconstruction for dendritic branching were set to 512X512, with an exposure rate of 1.6 per pixel, with one image acquired. All quantifications were performed in the somatosensory cortex, in sections of comparable rostro-caudal position. Dendrites and spines were quantified based on the mVenus fluorescence conveyed by the shRNA vector (pH1SCV2). Only dendrites that were largely parallel to the plane of the slice were analysed, 1 dendrite per neuron was analysed. Images were blindly acquired and analysed using Fiji^72^ (https://fiji.sc/). The morphometric analysis of dendritic spines along dendrites was analysed as previously described^32,56^ over a minimal dendritic length of 50 µm, starting from the object closest to the branchpoint with the main apical dendrite for apical dendrites or the soma for basal dendrites. The length of the dendritic segment was measured on the z projection. The reconstruction of dendritic branching was performed with the Fiji plugin Simple Neurite Tracer (SNT Neuroanatomy). Reconstructions were exported and analysed with a python script deposited on github (https://github.com/JosephZamith/FRMPD2B_project) to obtain branchpoint number along the dendrites. In vitro axon length was quantified on Fiji and sites along the length of the primary axon where secondary axonal branches arose were considered as branch points. Fluorescence intensity (FI) was used to quantify in vivo axonal branching, presynaptic boutons and promoter activity in our reporter assay as follows: FI = Integrated density/Area of interest – Mean grey value of background readings. To account for variation in the intensity of electroporation, FI of each area of interest was normalized to the FI of the electroporated area in L2/3 or to the FI of the pCAG tdTomato for the reporter assay.

### Whole transcripts RNA sequencing and analysis

Total RNA from human cerebral cortex was obtained from Takara (#636561). Library preparation and Nanopore sequencing were performed at the ENS Genomic core facility (Paris, France). Library preparation was done using SQK-LSK108 following manufacturer’s protocol (1D PCR Barcoding cDNA; ONT) optimized for cDNA sequencing. Briefly, 100 ng of total RNA was reverse transcribed for each sample, using custom polyT-VN and strand switching primers, with SuperScript IV (Life Technologies; 18090010), a quarter of the purified RT product was taken into PCR for amplification and barcodes addition (Barcoded primers form EXP-PBC001, ONT). 900 fmol cDNA libraries were loaded on an R9.4.1 flowcell after priming it according to the manufacturer’s protocol. Sequencing was performed with the standard 48-hour sequencGRAing protocol run on the MinION MkIB, using the MinKNOW software (versions 1.11.5). 3.78 million reads passing ONT quality filter (min quality set to 10) were obtained. Base-calling was performed by Guppy (version 6.4.2). The basic analyses were performed using the Eoulsan pipeline^73^, including read filtering, mapping, alignment filtering and read quantification. Poly N read tails were trimmed before mapping. Reads were then aligned against the human genome from Ensembl version 105 using Minimap2 (version 2.24)^74^ with the following options -x splice --eqx -- secondary=no --junc-bed human_bed12.bed. To compute gene expression, human genome annotation version 105 from Ensembl database was used. All overlapping regions between alignments and referenced exons (or genes) were counted and aggregated by genes using HTSeq-count 0.5.3^75^. Variant calling on *FRMPD2A* and *FRMPD2B* regions was performed by bcftools (version 1.10.2) (https://samtools.github.io/bcftools/bcftools.html). Isoforms analyses were performed using isoquant (v3.2.0)^76^. The analyses were guided on the human genome from Ensembl version 105, then the transcripts models were computed using human genome annotation version 105 with default options for ONT sequences. Isoquant was then used to annotate the genome without any annotation. Aligned transcripts reads were validated using Integrative Genomics Viewer (IGV)^77^.

### Transcription factor binding site analysis

Transcription factor binding site motifs were identified using the FIMO (Find Individual Motif Occurrences) tool, part of the MEME Suite^78^. To examine potential regulatory interactions, pFRMPD2B and phumPara sequences were scanned for occurrences of known transcription factor binding motifs sourced from JASPAR 2024 non-redundant vertebrate database, which contains position-specific weight matrices (PWMs) for 2,346 transcription factors. We used PsychEncode data retrieved from http://www.evolution.psychencode.org/ to filter out JASPAR 2024 based on TFs with cortical expression during synaptogenesis and set q-value to a maximum of 0.05.

### Statistics

Statistical analyses were performed with Prism 10 (GraphPad Software). For statistical analysis, normality of the distributions was assessed using the D’Agostino-Pearson normality test. In case of normal distributions, we used two-tailed Student’s t test or one-way analysis of variance followed by Tukey’s post hoc test. Non-normal distributions were assessed using the non-parametric Mann-Whitney test. In boxplots, whiskers correspond to the minimal and maximal values. Cumulative dendritic branchpoints were tested with a two-way repeated measures ANOVA. Other data represent mean ± SEM. Sample size and statistical tests are reported in each main and supplementary fig.. The significance threshold was placed at 0.05 (ns p > 0.05; ^*^p < 0.05, ^**^p <0.01; ^***^p<0.001).

## CODE AVAILABILITY

Original code used in this study is publicly available through GitHub: JosephZamith/FRMPD2B_project (https://github.com/JosephZamith/FRMPD2B_project). Code availability is specified in the Materials and Methods. Any additional information required to reanalyze the data reported in this paper is available from the lead contact upon request.

## ACKNOWLEDGMENTS

We thank all the members of the Charrier lab and M. Casado, M. Depp and R. Wimmer for discussion. We thank the acute transgenesis facility and the animal facility of IBENS for excellent support.

## AUTHOR CONTRIBUTIONS

Conceptualization: EC, JZ, CC. Methodology and investigation: global: EC, JZ, and CC; genomic analysis: FG, JZ, EC, CC; cortical organoids: PB, CB, MV, AB, CC; xenotransplantation: EC, MRV, WN, CC. Visualization: EC, JZ, CC. Funding acquisition: CC. Project administration: CC. Supervision: CC, AB. Writing original draft: EC, JZ, CC

## ADDITIONAL INFORMATION

Supplementary Fig. S1-7, Table S1.

