## Supplementary data for "The human-specific gene *FRMPD2B* couples postnatal experience to the expansion of cortical connectivity"

**The PDF file includes:**

Figs. S1 to S7

Table S1

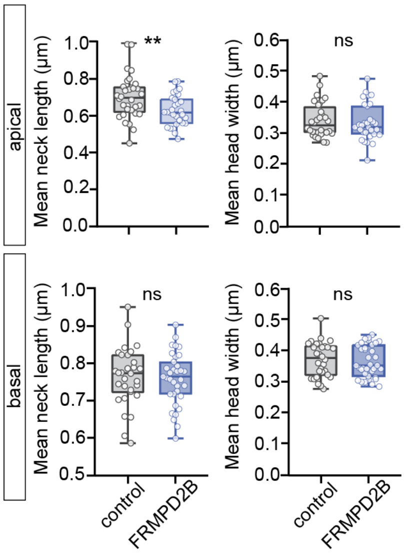


**Fig. S1 Morphometric analysis of dendritic spines.**
Quantification of mean neck length and head width in apical and basal dendrites of control and FRMPD2B-expressing neurons. Apical: $n_{control}$=32, $n_{FRMPD2B}$=31; basal: $n_{control}$=29, $n_{FRMPD2B}$=35 neurons. NS,*P>0.05*, ***P<0.01* by Mann-Whitney test.

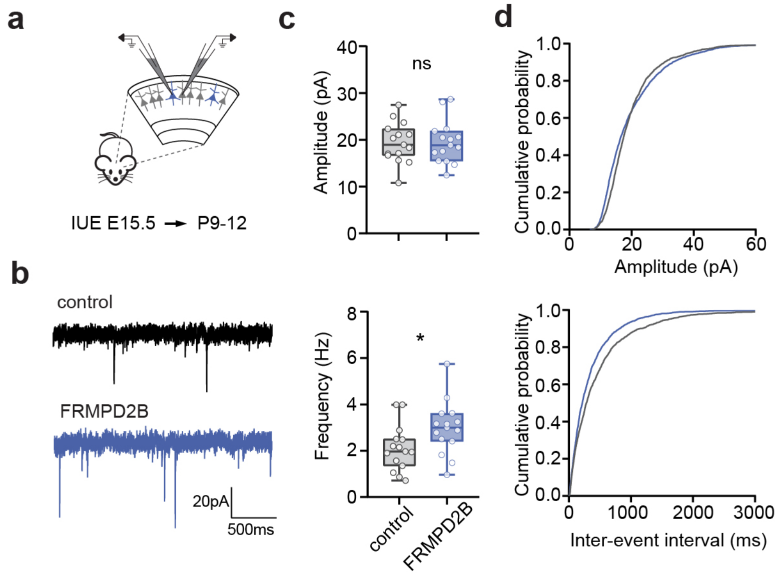


**Fig. S2 Increased number of active excitatory synapses in young mice following FRMPD2B expression.**
**a,** Whole-cell patch clamp recording in acute brain slices at P9-12 following sparse IUE at E15.5.
**b,** Representative mEPSC traces of control and FRMPD2B-expressing neurons.
**c,** Mean mEPSC amplitude and frequency, $n_{control}$=14, $n_{FRMPD2B}$=14 (6 mice).
**d,** Cumulative distribution of mEPSC amplitude and inter-event interval. NS,*P>0.05*, **P<0.05*, by Student t-test (c).

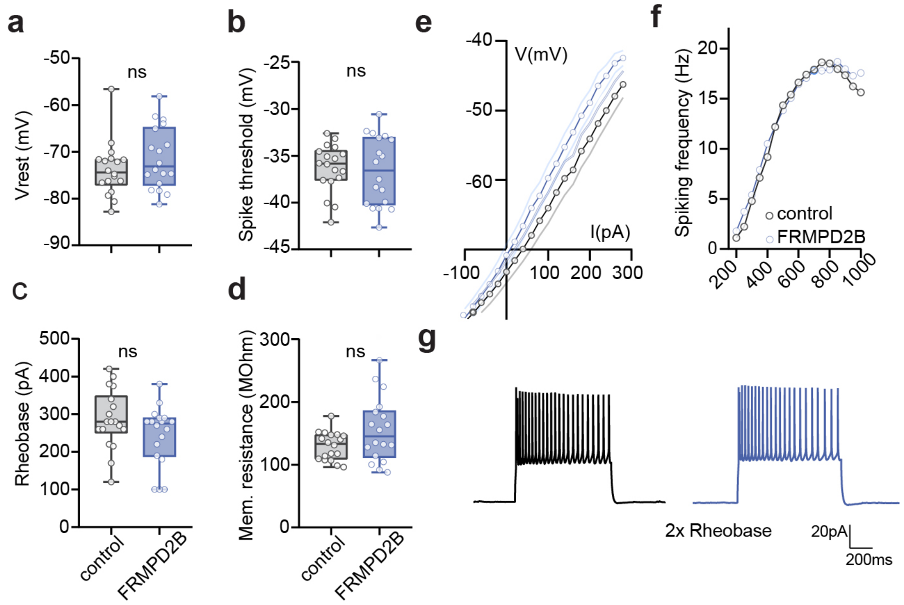


**Fig. S3 FRMPD2B does not alter L2/3 CPN intrinsic membrane properties.**

**a-e,** Quantification of intrinsic membrane properties in control and FRMPD2B-expressing neurons; $n_{control}$=20, $n_{FRMPD2B}$=20 neurons (7 brains per condition).

**f,** Action potential frequency in response to increasing steps of depolarizing current in control and FRMPD2B-expressing neurons.

**g,** Traces of action potentials evoked at twice the rheobase by 1s depolarizing current steps in whole cell current clamp recordings. NS,*P>0.05* by Students t-test.

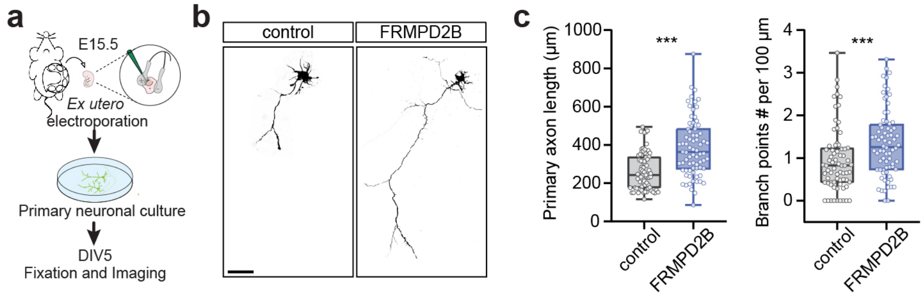


**Fig. S4 FRMPD2B enhances axonal length and branching of L2/3 CPNs *in vitro*.**
**a,** Experimental design: ex utero electroporation (EUE) was performed at E15.5. Neurons were dissociated and cultured for 5 days in vitro (DIV) before imaging.
**b,** Confocal images of cultured control and FRMPD2B-expressing L2/3 CPNs at DIV5. Scale bar: 50 µm.
**c,** Quantification of primary axon length and branching density (branch number per 100 µm) in control and FRMPD2B-expressing neurons, $n_{control}$=71; $n_{FRMPD2B}$=74 cells (from 4 mice per condition). ****P<0.001* by Mann-Whitney test.

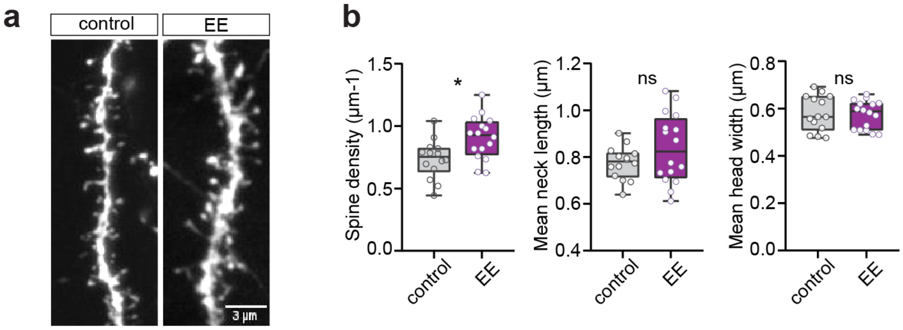


**Fig. S5 Increased spine density in basal dendrites of L2/3 CPNs in mice housed in enriched environment.**

**a,** Confocal images of basal dendrites from L2/3 CPNs at P14 expressing mVenus following IUE at E15.5 and housed in standing condition (control) or enriched environment (EE). Scale bar: 3 µm.

**b,** Mean spine density, mean neck length and head width in basal dendrites, $n_{control}$=14 neurons (5 mice), $n_{EE}$=16 neurons (7 mice). NS,*P>0.05*, **P<0.05* by Students t-test.

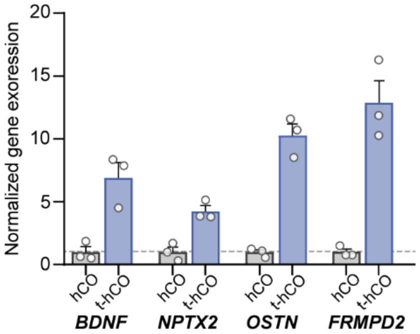


**Fig. S6 Expression of *FRMPD2* and other activity-regulated genes in human cortical neurons depending on xenotransplantation.**

Normalized expression of activity-regulated genes measured from glutamatergic neurons in human cortical organoids cultured for 8 months in vitro (hCO) or after 8 MPT into the rat cortex (t-hCO). Gene expression in counts per million was normalized to the mean expression of the non-transplantation group. Each dot represents a snRNAseq replicate. Note the 13-fold increase in *FRMPD2* expression in xenotransplanted neurons compared to neurons maintained in vitro. *OSTN*, which has gained MEF2-dependent regulation in primates[29],was also strongly upregulated after transplantation. Data extracted from[50].

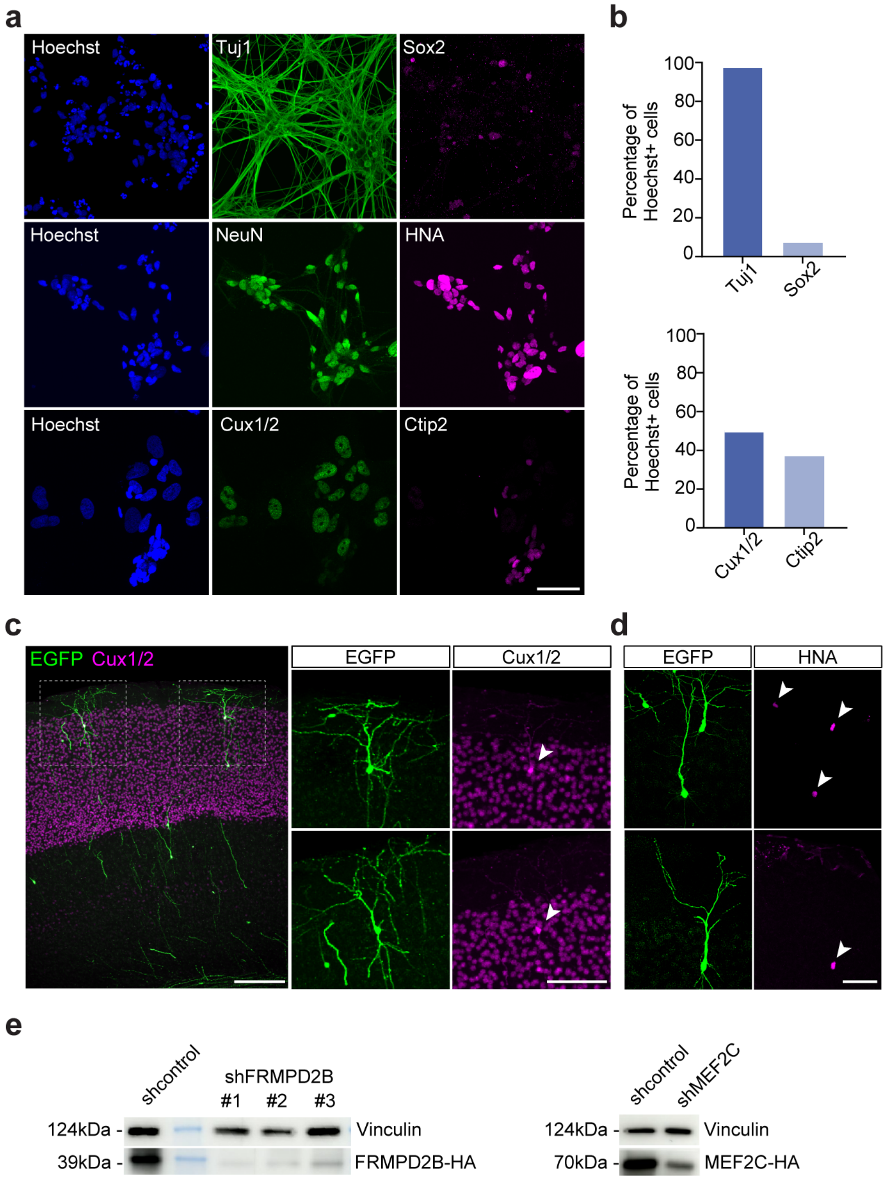


**Fig. S7 Characterization of the human cortical neurons and shRNAs used for xenotransplantation experiments.**

**a,** Characterization of human iPSC-derived cortical neurons after dissociation from cortical organoids and treatment with DAPT and ARA-C, as used for xenotransplantation experiments. Representative images showing expression of the neuronal markers Tuj1 and NeuN, the neural progenitor marker Sox2, the human-specific marker HNA, and the upper- and deep-layer cortical markers Cux1/2 and CTIP2. Nuclei were counterstained with Hoechst. Scale bar: 50 µm.

**b,** Quantification of neuronal differentiation and cell type identity.

**c-d,** Confocal images of EGFP-labelled xenotransplanted human neurons in the mouse cortex immunostained for Cux1/2 and HNA at 1 MPT. Arrowheads on the right highlight expression of Cux1/2 and HNA, confirming the upper-layer and human identity of xenotransplanted neurons. Scale bars: 100 µm (left); 50 µm (right).

**e,** Validation of shRNA constructs using Western blot analysis. Vinculin was used as a loading control.

| motif_id | motif_alt_id | start | stop | strand | score | p-value | q-value | matched_sequence |
|---|---|---|---|---|---|---|---|---|
| MA1596.1 | ZNF460 | -337 | -322 | + | 28.4831 | 9.80E-11 | 1.64E-07 | gcctcagcctcccgag |
| MA1596.1 | ZNF460 | -203 | -188 | + | 22.8764 | 1.65E-08 | 1.38E-05 | gcctcggcctcccaaa |
| MA1587.1 | ZNF135 | -202 | -189 | + | 19.0816 | 2.30E-07 | 0.000428 | cctcggcctcccaa |
| MA2121.1 | ZNF213 | -411 | -400 | + | 18.0894 | 2.59E-07 | 0.000428 | gcccaggctgga |
| MA1976.2 | ZNF320 | -670 | -651 | + | 16.8652 | 8.33E-07 | 0.00145 | ggaggacaagtgggcctcgg |
| MA1587.1 | ZNF135 | -336 | -323 | + | 16.0408 | 1.60E-06 | 0.00149 | cctcagcctcccga |
| MA0116.1 | Znf423 | -365 | -351 | + | 17.1124 | 1.68E-06 | 0.00184 | cgccccctggggttc |
| MA1587.1 | ZNF135 | -232 | -219 | + | 14.9286 | 2.96E-06 | 0.00311 | tctcgatctcctga |
| MA0685.2 | SP4 | -368 | -360 | - | 16.0674 | 3.34E-06 | 0.0038 | GGGGCGGAG |
| MA0079.5 | SP1 | -368 | -360 | - | 16.1327 | 3.34E-06 | 0.0049 | GGGGCGGAG |
| MA0146.3 | Zfx | -203 | -194 | + | 15.697 | 3.36E-06 | 0.0049 | gcctcggcct |
| MA0078.3 | Sox17 | -1021 | -1012 | - | 14.8239 | 3.36E-06 | 0.0049 | GGAACAATGG |
| MA1947.2 | ETV5::FOXO1 | -65 | -56 | + | 14.8429 | 4.03E-06 | 0.0049 | gaaaacagga |
| MA1596.1 | ZNF460 | -370 | -355 | + | 11.9663 | 6.82E-06 | 0.0049 | ggctccgccccctggg |
| MA1630.3 | ZNF281 | -368 | -359 | - | 13.573 | 7.26E-06 | 0.0049 | GGGGGCGGAG |
| MA1516.2 | KLF3 | -164 | -155 | + | 14.4255 | 7.67E-06 | 0.0049 | caccgcgccc |
| MA2331.1 | ZNF157 | -692 | -672 | - | 10.9242 | 7.87E-06 | 0.0049 | AGGGAAGCAGTATGTTTTCCC |
| MA0155.1 | INSM1 | -578 | -567 | + | 14.5102 | 8.11E-06 | 0.0049 | tggaaggggggtg |
| MA1522.2 | MAZ | -909 | -902 | - | 16.0816 | 8.13E-06 | 0.0049 | CCCCTCCC |
| MA1102.3 | CTCFL | -364 | -357 | - | 15.7245 | 8.13E-06 | 0.0049 | CAGGGGGC |
| MA1710.2 | ZNF257 | -342 | -333 | - | 14.5281 | 8.38E-06 | 0.0049 | GAGGCAGGAG |
| MA1930.2 | CTCF | -366 | -334 | - | 11.7982 | 8.83E-06 | 0.0049 | AGGCAGGAGAATGGCGTGAACCCCAGGGGGCGG |
| MA1712.2 | ZNF454 | -370 | -354 | + | 13.439 | 9.26E-06 | 0.00582 | ggctccgccccctgggg |
| MA1652.2 | ZKSCAN5 | -817 | -809 | + | 14.2844 | 9.34E-06 | 0.00594 | ggaggtgat |
| MA1653.2 | ZNF148 | -911 | -902 | - | 13.8163 | 9.86E-06 | 0.0061 | CCCCTCCCAG |
| MA0597.3 | THAP1 | -887 | -880 | - | 14.798 | 9.90E-06 | 0.00666 | GCAGGGCA |
| MA0597.3 | THAP1 | -802 | -795 | - | 14.798 | 9.90E-06 | 0.00799 | GCAGGGCA |
| MA0516.3 | SP2 | -368 | -360 | - | 14.427 | 1.03E-05 | 0.00877 | GGGGCGGAG |
| MA0139.2 | CTCF | -366 | -352 | - | 13.0164 | 1.07E-05 | 0.00877 | AACCCCAGGGGGCGG |
| MA1946.2 | ETV5::FOXI1 | -65 | -54 | + | 13.5606 | 1.15E-05 | 0.0128 | gaaaacaggagg |
| MA0742.2 | KLF12 | -368 | -360 | - | 13.9551 | 1.22E-05 | 0.0133 | GGGGCGGAG |
| MA0520.2 | Stat6 | -72 | -63 | - | 14.2241 | 1.32E-05 | 0.0137 | TTCCTGTGAA |
| MA1936.2 | ERF::FOXO1 | -65 | -54 | + | 13.4714 | 1.32E-05 | 0.0138 | gaaaacaggagg |
| MA1511.2 | KLF10 | -368 | -360 | - | 13.7978 | 1.40E-05 | 0.0139 | GGGGCGGAG |
| MA0496.4 | MAFK | -339 | -330 | + | 13.2061 | 1.40E-05 | 0.014 | ctgcctcagc |
| MA0497.2 | MEF2C | -277 | -267 | - | 13.1017 | 1.50E-05 | 0.0149 | CTAAAAATACA |
| MA0685.2 | SP4 | -910 | -902 | + | 13.8315 | 1.55E-05 | 0.0153 | tgggagggg |
| MA0071.1 | RORA | -454 | -445 | - | 13.7449 | 1.56E-05 | 0.0153 | AAAAAGGTCA |
| MA0787.1 | POU3F2 | -864 | -853 | + | 13.1774 | 1.61E-05 | 0.0154 | tcatgcaaatga |
| MA0116.1 | Znf423 | -364 | -350 | + | 9.11236 | 1.61E-05 | 0.0155 | gccccctggggttca |
| MA0746.3 | SP3 | -369 | -359 | + | 12.8654 | 1.62E-05 | 0.0156 | gctccgccccc |
| MA0105.4 | NFKB1 | -260 | -248 | + | 7.57692 | 1.63E-05 | 0.0168 | cggggtttcaccg |
| MA0627.3 | POU2F3 | -863 | -855 | + | 13.697 | 1.69E-05 | 0.0171 | catgcaaat |
| MA2122.1 | Ebf4 | -363 | -353 | - | 12.8085 | 1.69E-05 | 0.0174 | ACCCCAGGGGG |
| MA0087.3 | Sox5 | -1021 | -1014 | - | 14.3816 | 1.79E-05 | 0.0184 | AACAATGG |
| MA0077.2 | SOX9 | -1021 | -1014 | + | 12.939 | 1.79E-05 | 0.0184 | ccattgtt |
| MA0160.3 | NR4A2 | -454 | -447 | - | 14.2 | 1.79E-05 | 0.0202 | AAAGGTCA |
| MA1112.3 | NR4A1 | -454 | -447 | - | 14.6636 | 1.79E-05 | 0.0214 | AAAGGTCA |
| MA1728.2 | ZNF549 | -904 | -897 | - | 12.7154 | 1.80E-05 | 0.022 | GGCTGCCC |
| MA2099.1 | ZNF770 | -199 | -192 | + | 13.9541 | 1.80E-05 | 0.022 | cggcctcc |
| MA0052.5 | MEF2A | -276 | -267 | - | 13.1062 | 1.86E-05 | 0.0222 | CTAAAAATAC |
| MA0065.3 | Pparg::Rxra | -709 | -697 | - | 12.6306 | 1.96E-05 | 0.0228 | AGGCCAAAGAGCA |
| MA0773.1 | MEF2D | -277 | -266 | - | 10.6727 | 1.97E-05 | 0.0257 | ACTAAAAATACA |
| MA0516.3 | SP2 | -207 | -199 | - | 13.0899 | 2.03E-05 | 0.0262 | GAGGCGGGC |
| MA0741.1 | KLF16 | -369 | -359 | + | 12.7931 | 2.04E-05 | 0.0264 | gctccgccccc |
| MA1531.2 | NR1D1 | -221 | -208 | - | 9.94231 | 2.09E-05 | 0.0275 | GGATCACGAGGTCA |
| MA1564.2 | SP9 | -368 | -359 | + | 12.9273 | 2.10E-05 | 0.0275 | ctccgccccc |
| MA2341.1 | FEZF2 | -335 | -328 | + | 12.5571 | 2.20E-05 | 0.0275 | ctcagcct |
| MA0017.3 | NR2F1 | -225 | -214 | - | 13.3269 | 2.21E-05 | 0.0276 | CGAGGTCAGGAG |

| | | | | | | | | |
|---|---|---|---|---|---|---|---|---|
| MA1961.2 | PATZ1 | -911 | -901 | + | 12.382 | 2.22E-05 | 0.0276 | ctgggaggggc |
| MA0785.2 | POU2F1 | -863 | -855 | + | 12.25 | 2.41E-05 | 0.0276 | catgcaaat |
| MA0507.3 | POU2F2 | -864 | -852 | + | 10.75 | 2.49E-05 | 0.0286 | tcatgcaaatgat |
| MA0838.1 | CEBPG | -394 | -385 | + | 11.8163 | 2.61E-05 | 0.0286 | gtggcgcaat |
| MA0056.3 | MZF1 | -965 | -958 | - | 12.3577 | 2.67E-05 | 0.03 | CATCCCCA |
| MA2121.1 | ZNF213 | -207 | -196 | - | 12.187 | 2.75E-05 | 0.0309 | GCCGAGGCGGGC |
| MA1712.2 | ZNF454 | -899 | -883 | + | 11.4634 | 2.76E-05 | 0.0309 | gcctctaggccttgccc |
| MA1637.2 | EBF3 | -362 | -354 | - | 12.0325 | 2.79E-05 | 0.0311 | CCCCAGGGG |
| MA1953.2 | FOXO1::ELF1 | -65 | -53 | + | 12.1831 | 2.85E-05 | 0.0311 | gaaaacaggagga |
| MA0105.4 | NFKB1 | -260 | -248 | - | 5.40385 | 2.93E-05 | 0.0314 | CGGTGAAACCCCG |
| MA0159.1 | RARA::RXRA | -480 | -464 | - | 11.5102 | 2.96E-05 | 0.0317 | GGTTGACGCTGGGGACA |
| MA0753.3 | ZNF740 | -783 | -774 | - | 11.5169 | 2.97E-05 | 0.0318 | ACCCCCCAAC |
| MA1654.2 | ZNF16 | -605 | -585 | - | 6.09677 | 2.97E-05 | 0.0321 | AAGTGGGAGCTGAGCAAGGGC |
| MA1959.2 | KLF7 | -368 | -361 | - | 13.3371 | 3.11E-05 | 0.0321 | GGGCGGAG |
| MA1637.2 | EBF3 | -363 | -355 | - | 11.8455 | 3.16E-05 | 0.0328 | CCCAGGGGG |
| MA0017.3 | NR2F1 | -458 | -447 | - | 12.7885 | 3.18E-05 | 0.0338 | AAAGGTCAGGGA |
| MA2097.1 | ZNF75A | -835 | -824 | + | 10.9655 | 3.18E-05 | 0.0339 | acttttctcacc |
| MA2124.1 | Hmga1 | -566 | -559 | + | 13.2212 | 3.23E-05 | 0.034 | atttttat |
| MA1125.2 | ZNF384 | -449 | -442 | - | 13.5745 | 3.23E-05 | 0.035 | AAAAAAAA |
| MA1125.2 | ZNF384 | -448 | -441 | - | 13.5745 | 3.23E-05 | 0.0352 | AAAAAAAA |
| MA1125.2 | ZNF384 | -447 | -440 | - | 13.5745 | 3.23E-05 | 0.0353 | AAAAAAAA |
| MA1125.2 | ZNF384 | -446 | -439 | - | 13.5745 | 3.23E-05 | 0.0353 | AAAAAAAA |
| MA1125.2 | ZNF384 | -445 | -438 | - | 13.5745 | 3.23E-05 | 0.0354 | AAAAAAAA |
| MA1125.2 | ZNF384 | -444 | -437 | - | 13.5745 | 3.23E-05 | 0.0356 | AAAAAAAA |
| MA1125.2 | ZNF384 | -443 | -436 | - | 13.5745 | 3.23E-05 | 0.0357 | AAAAAAAA |
| MA1125.2 | ZNF384 | -442 | -435 | - | 13.5745 | 3.23E-05 | 0.0358 | AAAAAAAA |
| MA1125.2 | ZNF384 | -441 | -434 | - | 13.5745 | 3.23E-05 | 0.0358 | AAAAAAAA |
| MA1125.2 | ZNF384 | -440 | -433 | - | 13.5745 | 3.23E-05 | 0.0359 | AAAAAAAA |
| MA1125.2 | ZNF384 | -439 | -432 | - | 13.5745 | 3.23E-05 | 0.0359 | AAAAAAAA |
| MA1125.2 | ZNF384 | -438 | -431 | - | 13.5745 | 3.23E-05 | 0.0364 | AAAAAAAA |
| MA1125.2 | ZNF384 | -134 | -127 | + | 13.5745 | 3.23E-05 | 0.0364 | aaaaaaaa |
| MA1508.2 | IKZF1 | -70 | -63 | + | 11.813 | 3.26E-05 | 0.0369 | cacaggaa |
| MA1556.1 | RXRG | -356 | -343 | + | 9.89655 | 3.26E-05 | 0.0369 | gggttcacgccatt |
| MA2341.1 | FEZF2 | -407 | -400 | - | 12.2571 | 3.40E-05 | 0.0387 | TCCAGCCT |
| MA0815.1 | TFAP2C | -365 | -353 | - | 11.5862 | 3.49E-05 | 0.0393 | ACCCCAGGGGGCG |
| MA1727.2 | ZNF417 | -307 | -301 | + | 14.5506 | 3.60E-05 | 0.0394 | ggcgcca |
| MA0696.1 | ZIC1 | -365 | -352 | + | 8.21277 | 3.61E-05 | 0.0394 | cgccccctggggtt |
| MA1599.2 | ZNF682 | -370 | -360 | + | 11.2683 | 3.63E-05 | 0.0397 | ggctccgcccc |
| MA0786.2 | POU3F1 | -864 | -855 | + | 11.9157 | 3.64E-05 | 0.0433 | tcatgcaaat |
| MA1592.2 | ZNF274 | -961 | -950 | + | 10.5957 | 3.65E-05 | 0.0445 | gatgggtcctca |
| MA1562.2 | SOX14 | -1020 | -1012 | - | 12.3485 | 3.66E-05 | 0.0445 | GGAACAATG |
| MA0872.1 | TFAP2A | -365 | -353 | + | 11.7377 | 3.68E-05 | 0.0448 | cgccccctggggt |
| MA0814.3 | TFAP2C | -337 | -329 | - | 11.9187 | 3.71E-05 | 0.0448 | GGCTGAGGC |
| MA0742.2 | KLF12 | -910 | -902 | + | 11.2022 | 3.74E-05 | 0.0484 | tgggagggg |
| MA1954.2 | FOXO1::ELK1 | -65 | -53 | + | 11.2959 | 3.81E-05 | 0.0497 | gaaaacaggagga |

**Table S1 Transcription factor binding-site analysis for pFRMPD2B.**
Predicted transcription factor binding sites for the pFRMPD2B sequence, generated using FIMO (Find Individual Motif Occurrences; MEME Suite) with JASPAR vertebrate core motifs. The analysis was restricted to transcription factors (TFs) expressed in the cortex during the period of synaptogenesis. The table reports motif ID, TF name, start/stop positions, strand, score, p-value, q-value, and matched sequence.
